# Learning Gaussian Graphical Models from Correlated Data

**DOI:** 10.1101/2024.04.03.587948

**Authors:** Zeyuan Song, Sophia Gunn, Stefano Monti, Gina Marie Peloso, Ching-Ti Liu, Kathryn Lunetta, Paola Sebastiani

**Affiliations:** Institute for Clinical Research and Health Policy Studies, Tufts Medical Center, Boston, MA; The New York Genome Center, New York, NY, USA; Section of Computational Biomedicine, Boston University School of Medicine, Boston, MA 02218, USA; Bioinformatics Program, Boston University, Boston, MA 02215, USA; Department of Biostatistics, Boston University School of Public Health, Boston, MA; Tufts University School of Medicine, Boston MA; Data Intensive Study Program, Tufts University, Medford MA

**Keywords:** Gaussian Graphical Models, Corelated Data, Bootstrap, Polygenic Risk Score

## Abstract

Gaussian Graphical Models (GGM) have been widely used in biomedical research to explore complex relationships between many variables. There are well established procedures to build GGMs from a sample of independent and identical distributed observations. However, many studies include clustered and longitudinal data that result in correlated observations and ignoring this correlation among observations can lead to inflated Type I error. In this paper, we propose a Bootstrap algorithm to infer GGM from correlated data. We use extensive simulations of correlated data from family-based studies to show that the Bootstrap method does not inflate the Type I error while retaining statistical power compared to alternative solutions. We apply our method to learn the GGM that represents complex relations between 47 Polygenic Risk Scores generated using genome-wide genotype data from a family-based study known as the Long Life Family Study. By comparing it to the conventional methods that ignore within-cluster correlation, we show that our method controls the Type I error well in this real example.

## Introduction

In biomedical research it is often of interest to understand the network of complex relationships between various biological variables and other factors to improve disease diagnosis and prognosis, and to identify drug targets (Vamathevan et al., 2019). A Gaussian graphical model (GGM) is a powerful statistical machine learning tool that describes complex linear dependencies among many quantitative factors that are normally distributed using a Markov graph (Markowetz & Spang, 2007). The conventional method for learning a GGM is to perform hypothesis testing of the partial correlations that are derived from the normalized inverse of the variance-covariance matrix of the variables of interest (Whittaker, 2009).

The conventional method roots on the basic assumptions that the variables follow a multivariate normal distribution, and the sample data are independent and identically distributed observations. Unfortunately, the assumption of independent observations is violated in correlated data that arise from cluster sampling, or family-based studies. Talluri and Shete adapted the Lasso-penalized maximum likelihood estimator of the precision matrix by including the kinship matrix to account for the correlations introduced by family data (Talluri & Shete, 2014). However, the regression type analysis loses the computational advantages of the approach based on partial-correlation testing. Riberiro and Soler further leveraged the properties of family data for learning GGMs that are decomposed into the genetic and environmental networks (Ribeiro & Maria Pavan Soler, 2020). Their approach is interesting if the goal is to distinguish between genetic and non-genetic contributions to the association between the variables in the model. However, not all correlated data are from family-based studies.

In this work, we propose a Bootstrap algorithm to learn a GGM from correlated data. The advantage of this method is that there is no need to estimate the correlations within the clusters, and the approach is not limited to family-based data. We show through a comprehensive simulation study that our algorithm controls the Type I error well, while retaining good statistical power. We also apply our method in a real-world example to show the dramatic impact of ignoring correlated data when building a GGM.

## Methods

### Background: Learning Gaussian Graphic Models from Independent Observations

A Gaussian Graphic Model (GGM) is a statistical model that represents properties of marginal and conditional independencies of a multivariate Gaussian distribution using an undirected Markov graph(Lauritzen, 1996; Whittaker, 2009). The key rule of an undirected Markov graph is that two variables are conditionally independent given all the other variables in the graph if they are not connected by an edge. Let ***Y*** = (*Y*_1_, *Y*_2_, *Y*_3_, …, *Y*_*p*_)^*T*^ be a p-dimensional random vector with a multivariate normal distribution with mean vector µ and covariance matrix *Σ*:

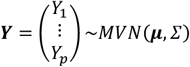

Let *G* denote the associated undirected Markov graph from the set (*V, E*) where *V* = {1,2, …, p} is the vertex set corresponding to the univariate random variables in ***Y***, and the edge set *E* = {*E*_*i,j*_ : *i, j* ∈ *V, i* ≠ *j*} describes the conditional dependency of random variables in ***Y*** (Kolaczyk, 2009). The strength of the conditional dependency of *Y*_*i*_ and *Y*_*j*_ given all other variables in ***Y*** is measured by the partial correlation ρ_*ij*_ that is defined as:

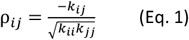

where *k*_*ij*_ is the (*i, j*)^*th*^ entry of the precision matrix *K* = *Σ*^−1^ (Whittaker, 2009). An edge exists between two vertices if the partial correlation between the two Gaussian random variables is not 0, i.e.

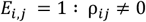

Figure 1 presents an illustrative GGM depicting the partial correlation network of a vector ***Y*** = (*Y*_1_, *Y*_2_, *Y*_3_, *Y*_4_)^*T*^. The Markov graph shows that *Y*_2_ is independent of *Y*_3_ and *Y*_4_ conditioned on *Y*_1_ (d-separation), and this relationship is represented by the missing edges *E*_2,3_ = 0 : ρ_23_ = 0 and *E*_2,4_ = 0 : ρ_24_ = 0. *Y*_1_ and *Y*_3_ are dependent on each other when conditioned on *Y*_2_ and *Y*_4_, which can be described by the existing edge *E*_1,3_ = 1 : ρ_13_ ≠ 0.

**Figure 1.**
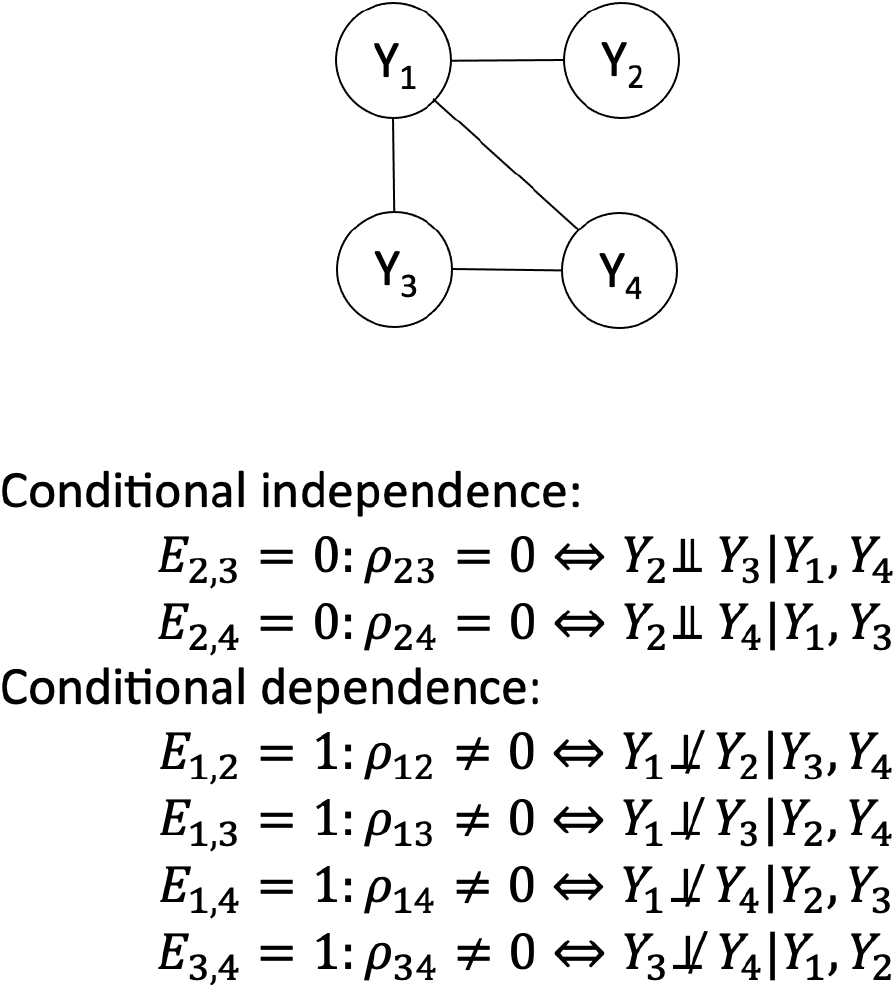
Example of a GGM with 4 vertices and 4 edges.

In general, the null hypothesis of conditional independence of *Y*_*i*_ and *Y*_*j*_ given all the other variables, say *H*_0_: ρ_*ij*_ = 0 against the alternative hypothesis *H*_1_: ρ_*ij*_ ≠ 0, can be tested using the Fisher’s z-transformation test:

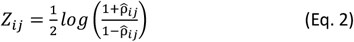

where 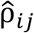 is the estimate from a sample with sample size *n* (Fisher, 1921). The distribution of the statistic under the null hypothesis *H*_0_: ρ_*ij*_ ≠ 0 can be approximated by

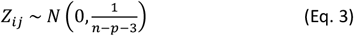

### Clustered Data and Issues

The statistical methods that are typically used to learn the structure and the parameters of a GGM from a sample data set assume that the observations are independent and identically distributed. However, the assumption of independence is violated in studies that collected correlated data such as cluster-based sampling or family-based recruitment (Laird, 2004). Subjects within a cluster are correlated due to shared environment components or sharing of genetic factors in family-based studies (Wojczynski et al., 2022). Failure to account for these correlations can lead to biased estimates and false positive results (Cannon et al., 2001).

In the analysis of cluster data, often an exchangeable covariance structure is assumed, where the correlation of pairs of subjects in the same cluster is constant. Family data is a special type of clustered data in which each family is a cluster unit. The correlation structure of family data can be more complex with correlations between pairs of subjects that can be determined by linked to their family relationship and shared environment. In genetic studies, the variance of a trait is commonly decomposed into two components: the independent environmental and the correlated genetic components (Almasy & Blangero, 1998; Amos, 1994). Denote by *y*_*ij*_ the observation of a variable *Y* in the *j*^*th*^ individual from the *i*^*th*^ family, the effect of the two components of variance can be parameterized as

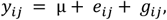

where 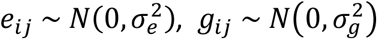, and the environmental factors *e*_*ij*_ are assumed to be independent while the genetical components *g*_*ij*_ are independent between different families and dependent within families. Therefore, for any two subjects from different families, the observations *y*_*ij*_ and *y*_*kl*_ are independent of each other.

For two subjects from the same family, *y*_*ij*_ and *y*_*ik*_ are dependent due to the covariance introduced by *g*_*ij*_ and *g*_*ik*_:

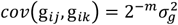

where *m* describes the degree of relatedness between the two individuals. The generating model for one trait with sample data 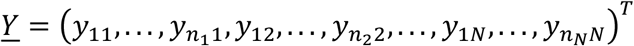 can be written as

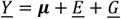

where 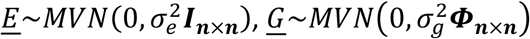, *n*_*i*_ is the number of subjects in the *i*^*th*^ family, *N* is the total number of families and 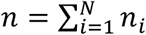. Here ***Φ***_***n*×*n***_ is a block diagonal matrix called the relatedness matrix. The elements *i, j* of ***Φ***_***n*×*n***_ represent the relatedness between individuals *i, j* and are 0 when *i, j* are from different families, and they are 2^−*m*^ when *i,j* are a *m*-*th* degree relative pair (Lange, 2022). Observations from family data are correlated because of this shared genetic component 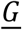. Ignoring this correlation could inflate the detection of false association between variables and introduce false edges in the GGM.

### Bootstrap Algorithm on Clustered Data

The Bootstrap method, introduced by Efron (Efron, 1979), is a widely used resampling technique for statistical inference and hypothesis testing. It involves generating new samples by sampling the data with replacement and then using these samples to estimate the distribution of a statistic of interest. Sherma and Cessie suggested that the Bootstrap method could also be used to address issues with correlated data by resampling clusters instead of individuals (Sherman & Cessie, 1997), and Borecki and Province introduced a family-based bootstrap approach in which the sample units are families and familial relations are ignored in the estimation phase (Borecki & Province, 2008).

Here we propose a generalization of the family-based Bootstrap algorithm introduced by Boreki and Province to learn GGMs that account for correlated observations. The steps of the proposed Bootstrap algorithm are as follows:

1. For *t*=1,2,…, *T*
  i. draw *c*% of clusters with replacement from the cluster data, e.g. draw 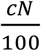 families with replacement;
  ii. estimate the variance-covariance matrix *Σ*^(*t*)^ from the data resampled in 1) and calculate the partial correlation matrix *P*^(*t*)^ using *Σ*^(*t*)^ at this *t*^*th*^ iteration using the equation:

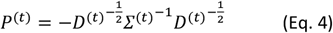

where *D*^(*t*)^ is a diagonal matrix from the diagonal elements of 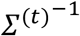;
  iii. for each pair of variables *Y*_*i*_ and *Y*_*j*_, calculate the Fisher’s transformation statistics 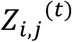 using (Eq. 2).
2. Repeat i)-iii) steps N times and build the standardized *Z* statistics as:

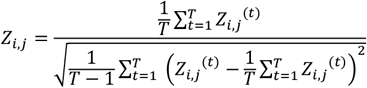

That is approximately normally distributed as *N*(0,1).
3. The null hypothesis is rejected at a given test size α if |*Z*_*i,j*_| > *Z*_α/2_ and the confidence interval of 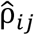 can be obtained as

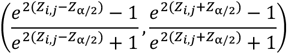
4. Obtain estimates of the precision matrix 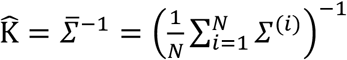 and thereby calculate the estimates of the partial correlation matrix 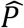 using (Eq. 4) with 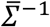.

### Simulation studies

#### Family Data Simulation

It is very challenging to simulate a data set of correlated variables with also correlated observations. Therefore, we describe in detail how we simulated the data for this evaluation. We begin with the generating model for a single trait *Y*:

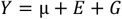

where 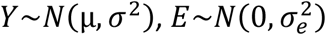 and 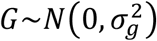 and 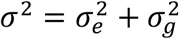.

We consider a simple scenario with two parents and one offspring, and we denote by *Y*_*f*_, *Y*_*m*_ and *Y*_*o*_ the value of *Y* in the father, mother, and offspring respectively. The parent-offspring pair is a 1^st^ degree relative pair and the genetic covariance between each parent and offspring is well known to be 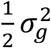 (Lange, 2022). Using this information, the distribution of the vector ***Y*** = (*Y*_*f*_, *Y*_*m*_, *Y*_*o*_)^***T***^ is

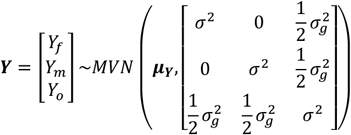

and we can use the derivation of the conditional distribution from a Multivariate normal distribution (Majumdar & Majumdar, 2019) to derive the distribution of *Y*_*o*_ conditionally on their parents’ data as:

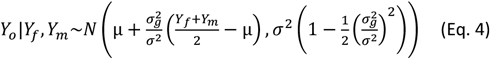

Note that this distribution depends on the ratio 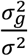 that is known as the heritability *h*^2^:

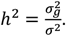

Suppose now that we have *p* variables 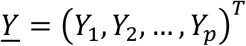 and we denote as 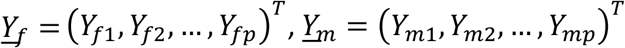 and 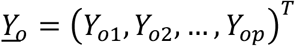 the data for father, mother, and offspring respectively. The joint distribution of 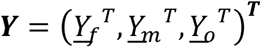 is

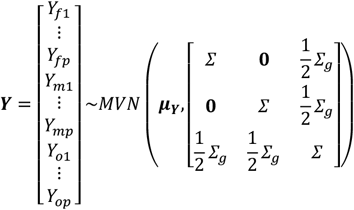

where *Σ* is the *p* × *p* variance-covariance matrix of the random vector (*Y*_1_, *Y*_2_, …, *Y*_*p*_)^*T*^, and *Σ*_*g*_ is the *p* × *p* variance-covariance matrix introduced by the genetic correlation. The inverse of the variance covariance matrix *Σ* is important here since the zero-entries of the off-diagonal terms represent the conditional independence among the variables *Y*_*i*_ and *Y*_*j*_ represented by the edge *E*_*i,j*_ = 0 in the GGM network. The conditional distribution of 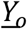 given their parents is derived as detailed in supplemental materials:

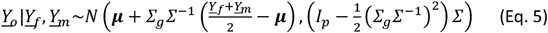

In Equation (5), *I*_*p*_ is the *p* × *p* identity matrix. Here we define the *p* × *p* heritability matrix *H* as:

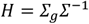

The diagonal elements of *H* represent the heritability of each trait while the off-diagonal values measure the genetic correlations between traits. If the traits are genetically independent of each other, *H* would be a diagonal matrix. Denote by ⊗ the Kronecker product. The variance-covariance matrix of ***Y*** can also written as:

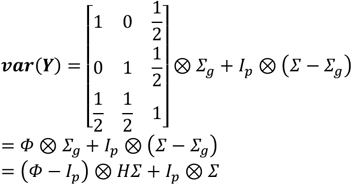

By the derivation above we obtained the formula for simulating multiple traits from families with parents and offspring structure:

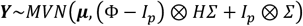

where we are interested to learn the GGM network with respect to *K* = *Σ*^−1^.

#### Simulation Settings

In the simulation study, we set *H* = *h*^2^*I*_*p*_ to generate genetically independent traits with different degrees of heritability. We simulated a mixed family structure where half of the families consisted of two parents and one offspring, while the other half consisted of two parents and five offspring. We simulated observations assuming three different numbers of families: 40, 120, and 360 families, resulting in sample sizes of n = 200, 600, and 1800, respectively. We varied the heritability values *h*^2^ from 0 to 0.95 with increments of 0.125, 0.25, 0.5, and 0.75. We also generated 1000 datasets for each combination of sample size and heritability. We simulated data from three different GGMs with Markov graphs depicted in Figure 2. The first model included three variables that were marginally independent of each other, so that the Markov graph did not include any edge (Figure 2a). The second model was represented by a chain graph describing two variables *Y*_1_ and *Y*_3_ conditionally independent given *Y*_2_ (Figure 2b). The third model was a triangle tail graph that described *Y*_1_, *Y*_2_ and *Y*_3_ connected to each other and *Y*_4_ is independent of *Y*_1_ and *Y*_2_ conditional on *Y*_3_ (Figure 2c). The *p* × *p* variance covariance matrix *Σ*, precision matrix *K*, and corresponding partial correlation matrix *P* for each graph are displayed in Figure 2. Without loss of generality, we set the mean µ of each variable to be 0.

**Figure 2.**
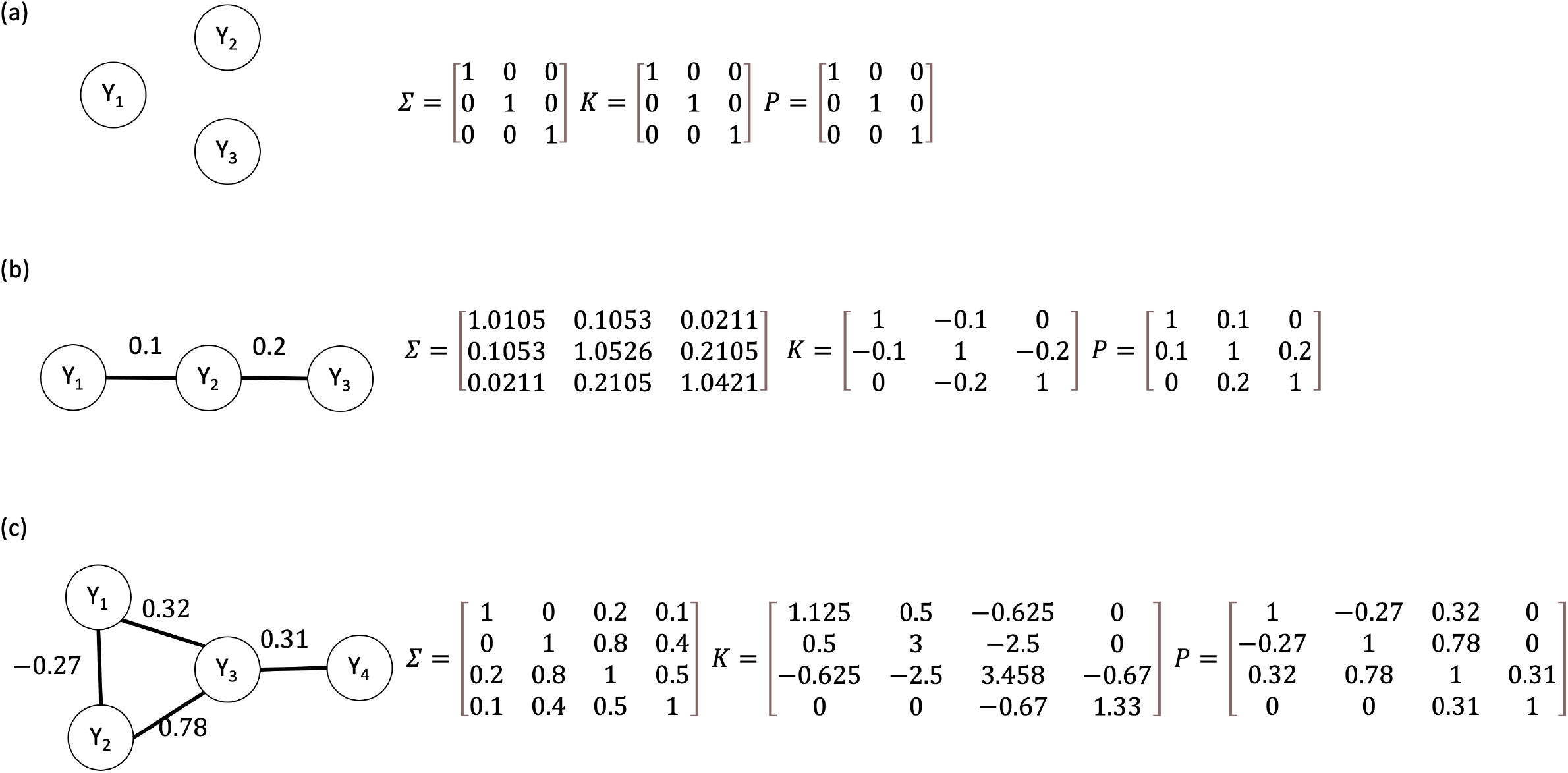
(a) the independence graph (b) the chain graph (c) the triangle tail graph Markov models that are used in the simulation studies.

In each scenario, we learned the GGM structure by using:

- The Fisher’s transformation test for partial correlations ignoring the family-based design.
- Our proposed cluster-based bootstrap algorithm, in which we resampled 50 and 200 datasets, and for each resample, we drew 50%, 75% and 100% of families with replacement.

We used a level of statistical significance *α* = 0.05 for the Hypothesis test of each *ρ*_*i,j*_. The FPR was estimated by the proportion of incorrect edges found to be significant. The power of the Hypothesis tests was evaluated using this algorithm: when inflation in FPR was presented in the conditionally independent variable pairs in one graph, we corrected the level of significance *α* by mean inflation rate, which is the average of the FPR divided by *α*, and conducted a new hypothesis test at *α*^∗^:

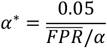

We repeated this process until *α*^∗^ = 0.05 and then used this adjusted significance level to estimate the power as the proportion of edges found between the variable pairs that were connected in the data model.

### Simulation Results

#### Inference

Using 1000 simulations, we obtained estimates of the false positive rate (FPR) by examining the proportions of edges between variables whose partial correlations are 0 in the three GGMs. Figure 3 summarizes the results of the FPR for different scenarios and methods. The FPR of the Fisher’s test that ignores the family structure increases across all three graphs as the heritability levels increase and the inflation rates increase by 2-4 fold as heritability exceeds 0.5. When the number of families was as small as 40, the Bootstrap algorithm that samples 100% of families exhibits a small but inflated FPR of approximately 1.5-fold across varying heritability levels. However, as the number of families increases to 120 and 360, sampling 100% of families consistently maintains the FPR at 0.05 across all heritability levels. Notably, the Bootstrap algorithm that samples 75% of families also maintains the FPR at 0.05 when the number of families is 40, but as the number of families increased, it consistently deflated the FRP. Similarly, the Bootstrap algorithm that samples 50% of families always suffered of deflated FRP. In summary, when the number of families exceeds 120, the Bootstrap algorithm that samples 100% of families would consistently maintain the FDR at 0.05 at varying heritability levels.

**Figure 3.**
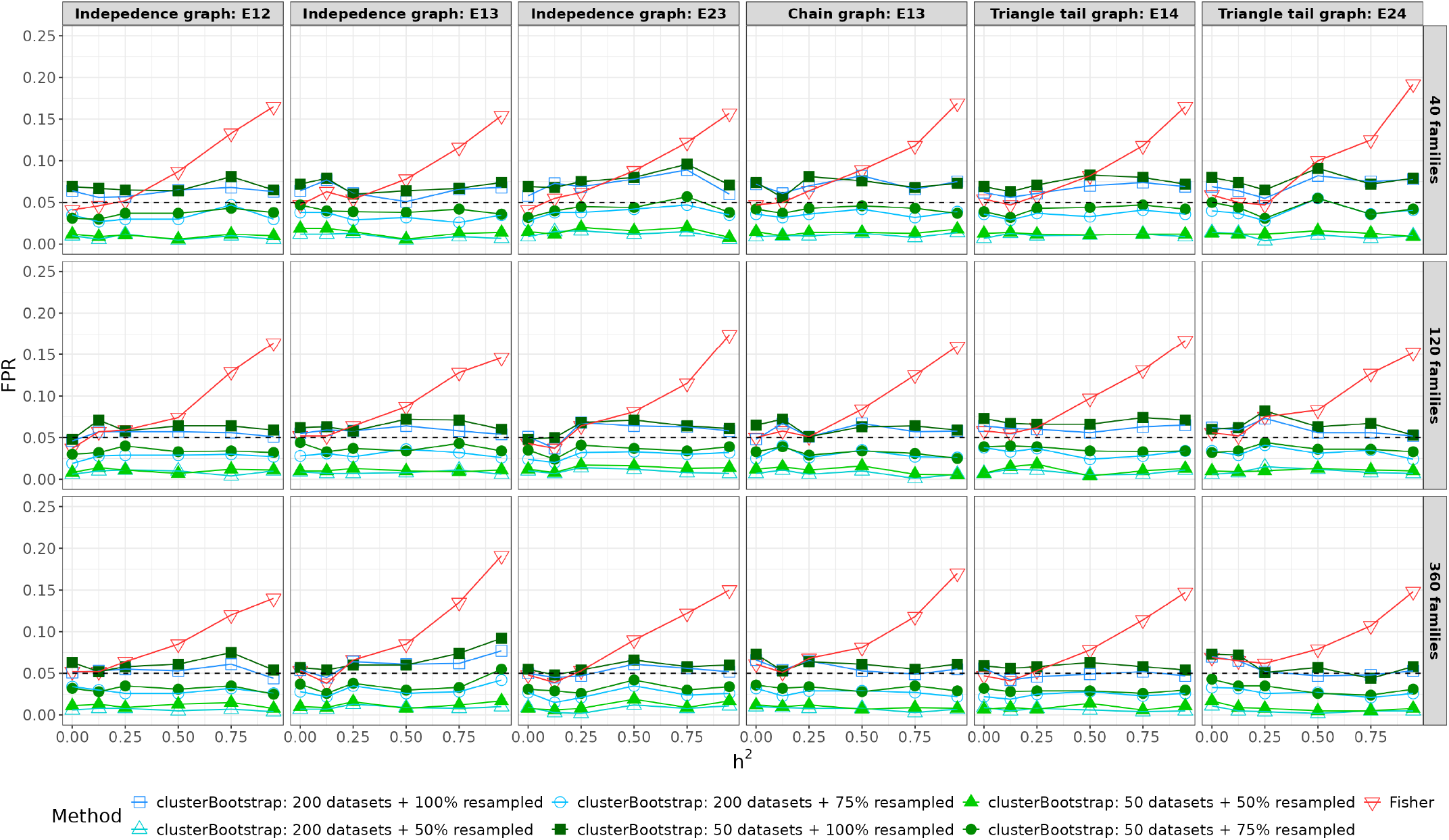
False Positive Rates at *α* = 0.05 across varying heritability for each *E*_*i,j*_ = 0 in all three graphs using Bootstrap (clusterBootstrap) and Fisher’s methods (Fisher).

Figure 4 summarizes the statistical power of detecting true edges using both the Fisher’s test and the proposed Bootstrap algorithm as a function of the heritability. Notably, as the heritability increases, the statistical power decreases particularly when sample sizes are small and partial correlations between variables are close to 0. However, when the number of families exceeds 360, the power remains consistently above 0.8, irrespective of the heritability values ranging from 0 to 0.95. For smaller family sizes of 40, a power above 0.8 was achieved as long as the magnitude of the partial correlations exceeded 0.3, regardless of heritability.

**Figure 4.**
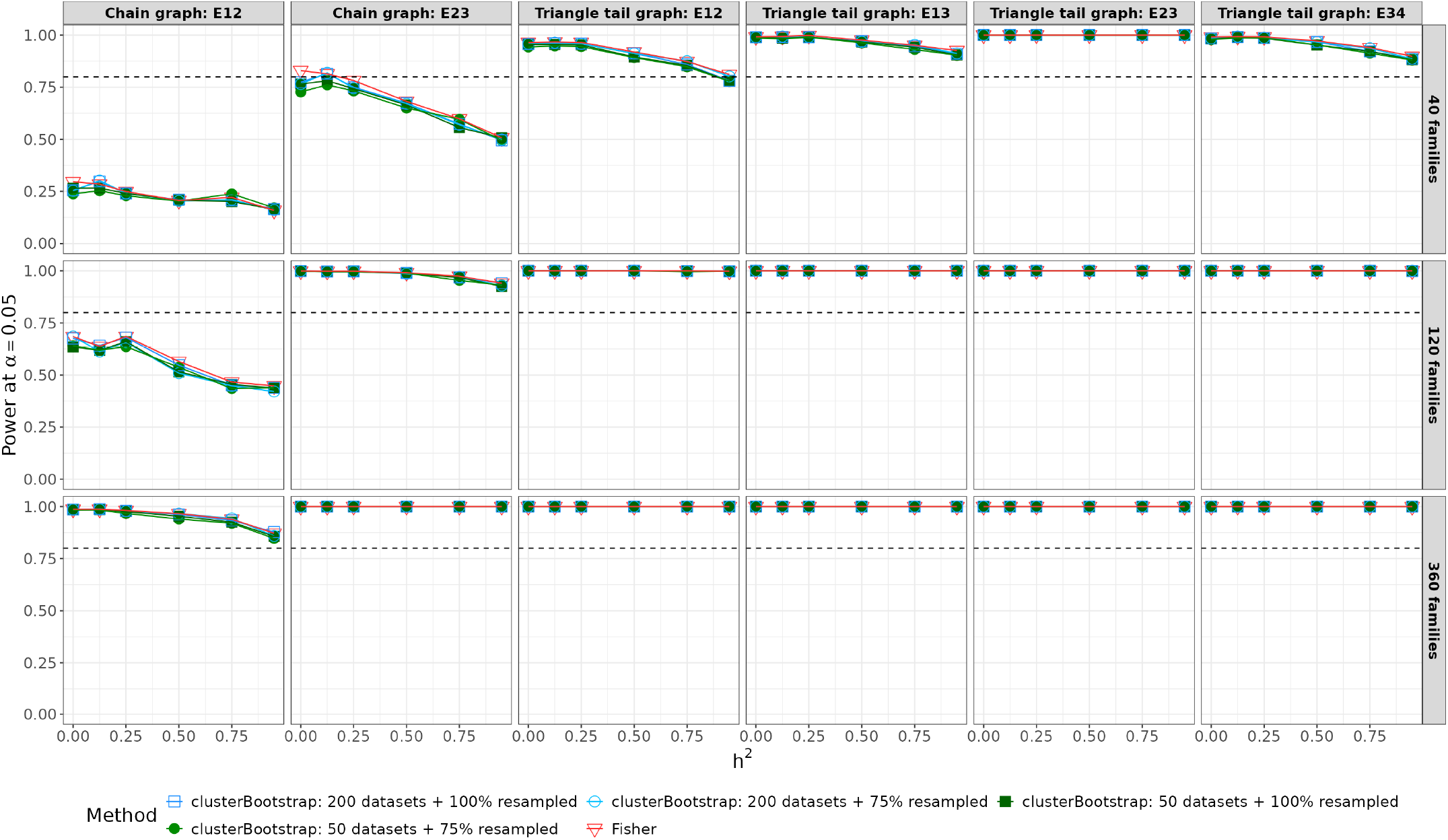
Power at *α*^∗^= 0.05 across varying heritability for each *E*_*i,j*_ = 1 for all three graphs using Bootstrap (clusterBootstrap) and Fisher’s methods (Fisher). *α*^∗^ is the adjusted significance level as described in the power evaluation section.

Comparing the proposed Bootstrap algorithm with the Fisher’s test, we observed comparable power across all scenarios when we controlled the Type I error rate. Our results indicated that bootstrapping 50/200 datasets with 100% families resampled yielded the best performance in terms of both power and FPR. However, reducing the proportion of families resampled may lead to an over-correction of the FPR.

#### Estimation

Figure 5 presents the estimates of the partial correlations between *Y*_1_ and *Y*_3_ in the triangle-tail graph. Both the naïve partial correlation estimates ignoring the correlated data nature and the Bootstrapped partial correlation estimates are unbiased across varying heritability and numbers of families [Supplemental Figure 1]. With increasing sample sizes, the standard deviations of these estimates narrow down as expected. This result indicates that the estimation can be performed using either method.

**Figure 5.**
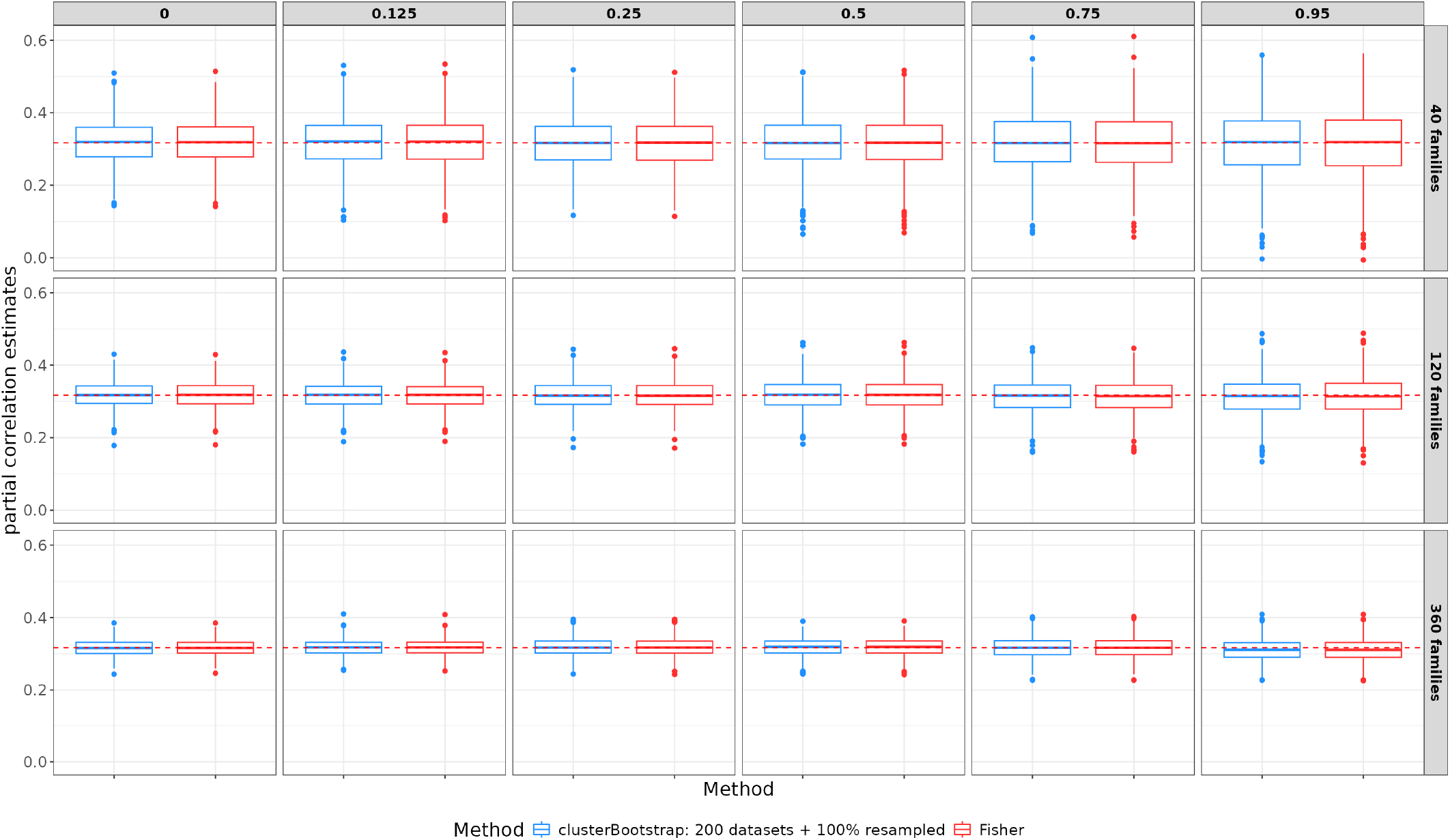
Estimates of the partial correlation at varying heritability (0-0.95) and numbers of families for *E*_1,3_ in the triangle tail graph using Bootstrap and Fisher’s methods. Red dash line reflects the true value.

### Application to Modeling the Dependency of Polygenic Risk Scores in a Family-Based Study

#### Study Cohort

We applied this new algorithm to characterize the mutual correlations between 47 polygenic risk scores in the Long Life Family Study (LLFS).

#### Data

LLFS is a family-based study of healthy aging and longevity that recruited over 5000 family members in approximately 550 families selected for familiar longevity. Participants were enrolled at three American field centers in Boston, Pittsburgh, and New York, along with a European field center based in Denmark (Sebastiani et al., 2009; Wojczynski et al., 2022). Socio-demographic, medical history data, current medications, physical and cognitive function data, and blood samples were collected via in-person visits and phone questionnaires for all subjects at the time of enrollment and during follow-ups (Elo et al., 2013; Newman et al., 2011). Genome-wide genotype data were generated at the Center for Inherited Disease Research (CIDR) using the Illumina Omni 2.5 platform, and genotype calls were cleaned as described in (Bae et al., 2013). The genotype data were imputed with Michigan Imputation Server to the HRC panel (version r1.1 2016) (Das et al., 2016). Genome-wide genotype data are available from dbGaP (dbGaP Study Accession: phs000397.v1.p1). We augmented the genetic data in the LLFS using approximately 3500 samples that we used as controls in other studies of longevity (Bae et al., 2013). The genotype data are accessible through the specified protocol available at http://www.illumina.com/documents/icontroldb/document_purpose.pdf. The genome-wide genotype data were generated employing several Illumina SNP arrays, with quality control performed as described in (Sebastiani et al., 2012), and were imputed the same way as the LLFS cohort.

#### Learning Polygenic Risk Score Network

Polygenic Risk Scores (PRS) for 54 health outcomes using genetic data of 8,190 samples were calculated as reported by (Gunn et al., 2022). We further cleaned the PRS data by removing two PRS with very skewed distribution [Supplemental Figure 2] and additional five PRS that had several potential outliers that lie 4 standard deviations away from the means [Supplemental Figure 3]. We learned the partial correlation networks of the remaining 47 PRS using three methods: Fisher’s transformation test with independent subsets of the data yielding a sample size of N=4193, Fisher’s test on all data ignoring the correlation within families (N=8190), and the proposed bootstrap algorithm (N=8190). In the first method, we generated independent subsets by randomly sampling one subject per family. In the second method, we applied Fisher’s test to all data. With the Bootstrap algorithm we sampled 1000 datasets with 100% of families sampled each sampling. We applied Bonferroni correction to control the family-wise error rate (FWER) to be 0.05.

#### PRS Network Comparison

Figure 6 displays the networks constructed using the three methods. The Fisher’s test on independent subsets identified 85 edges among 4193 independent subjects, while ignoring the correlations within families identified 143 edges using data from 8190 subjects. The Bootstrap method applied to the data set of 8190 subjects identified a total of 108 edges (Table 1 and Figure 7). As expected, this number was between the previous two methods as the analysis of only the independent observations is less powerful since the sample size is reduced by almost 50%, while the method that analyzes all the data ignoring correlations within families likely introduced false positive edges. Table 1 and Figure 7 showed that the three algorithms identified 78 edges in common. The Bootstrap algorithm identified additional 30 more edges that were also identified with the Fisher’s test applied to all samples. However, the latter method identified additional 30 edges that are not identified by neither the Fisher’s on independent sample nor the Bootstrap algorithm.

**Table 1.** Comparison of numbers of common edges inferred by the three methods.

| Edges by Bootstrap = 0 |  | Edges by Fisher's (all sample) |  |
| --- | --- | --- | --- |
|  |  | 0 | 1 |
| Edges by Fisher's (iid sample) | 0 | 936 | 30 |
|  | 1 | 2 | 5 |

| Edges by Bootstrap = 1 |  | Edges by Fisher's (all sample) |  |
| --- | --- | --- | --- |
|  |  | 0 | 1 |
| Edges by Fisher's (iid sample) | 0 | 0 | 30 |
|  | 1 | 0 | 78 |

**Figure 6.**
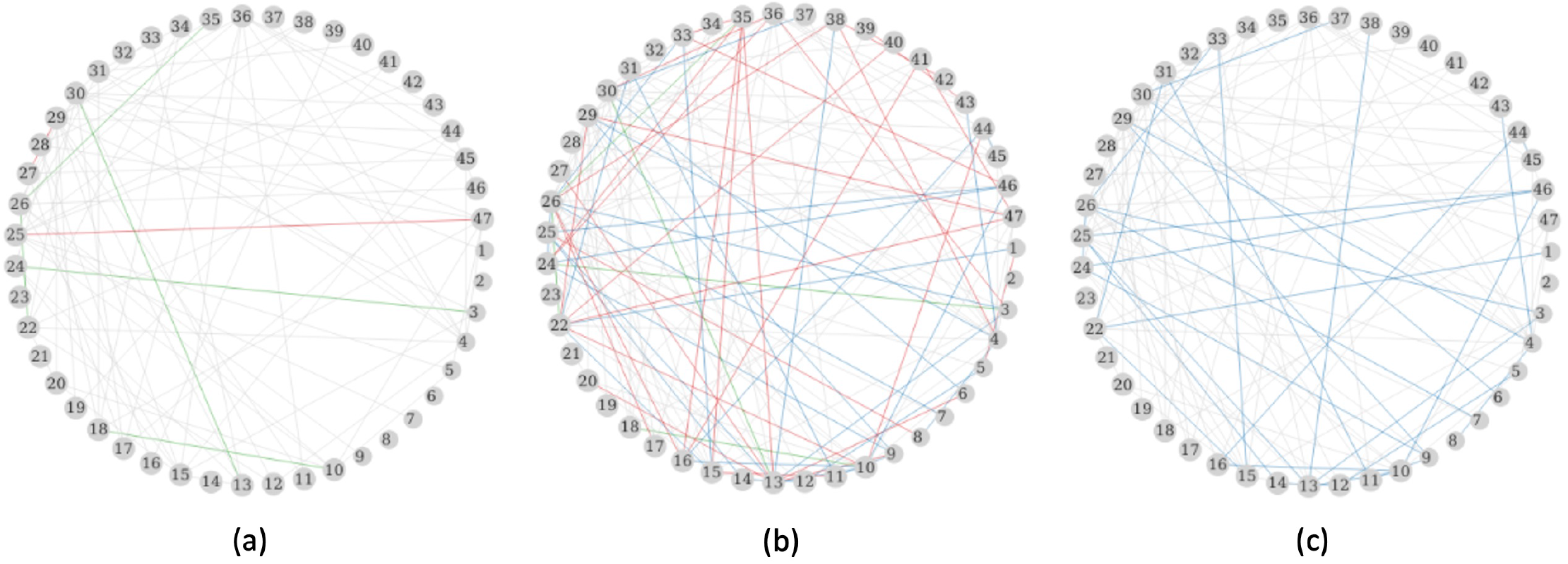
PRS networks inferred with Bonferroni correction using (a) Fisher’s test on sampled iid data, (b) Fisher’s test on all data, and (c) Bootstrap resampling 1000 datasets and 100% families. Grey edges are edges identified in all three approaches; green edges are intersections by (a) and (b); blue edges are intersections by (b) and (c); red edges are identified only by the corresponding approach. The numbers in each circle is a PRS as listed in **Supplementary Table 1**.

**Figure 7.**
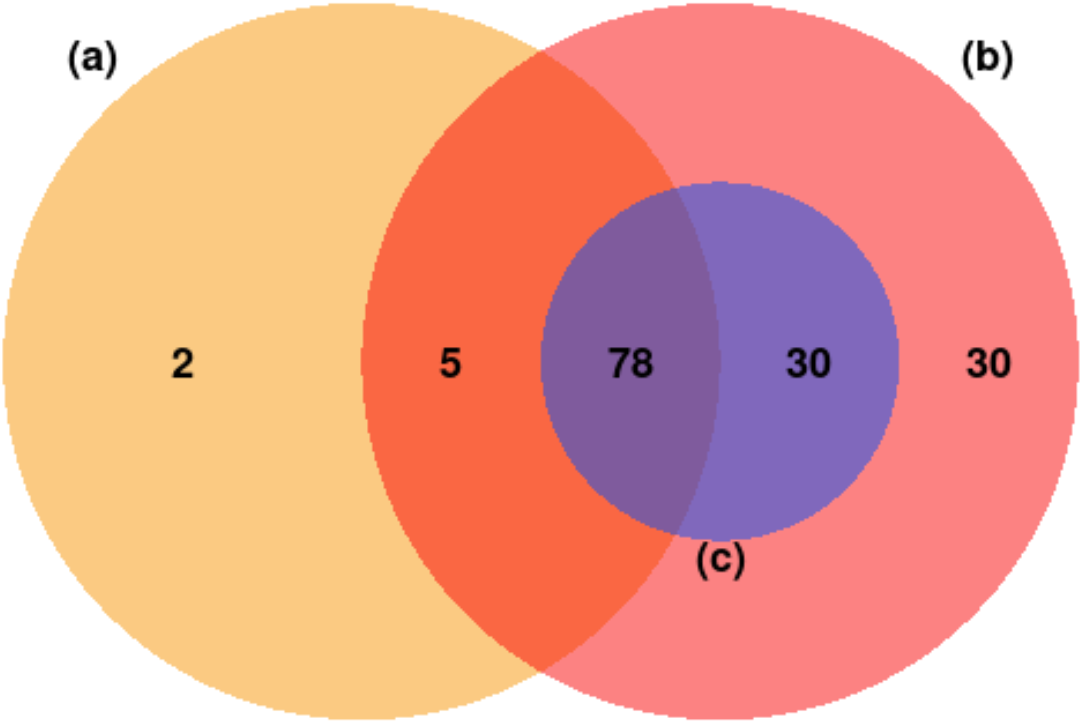
Venn diagram of common edges of the PRS networks inferred with Bonferroni correction using (a) Fisher’s test on sampled iid data, (b) Fisher’s test on all data, and (c) Bootstrap resampling 1000 datasets and 100% families.

#### Evaluation of Computation Time

We evaluated the computation time of the Bootstrap algorithm by calculating the CPU time of the resampling steps and the inference steps. We ran the evaluation using a single computer node with 1 core and R version 4.1.1. We sampled 40, 400, and 4000 families from the LLFS data, and sampled 10, 20, 40 PRS. For each scenario, we obtained the computational time for 50, 200 and 1000 iterations with 100% of the families resampled each iteration. The algorithm finished in 65 seconds in the scenario with 40 PRS, 4000 families and 1000 iterations as shown in Figure 8. Notably, the resampling step took 52 seconds that makes up to 81% of the total CPU time.

**Figure 8.**
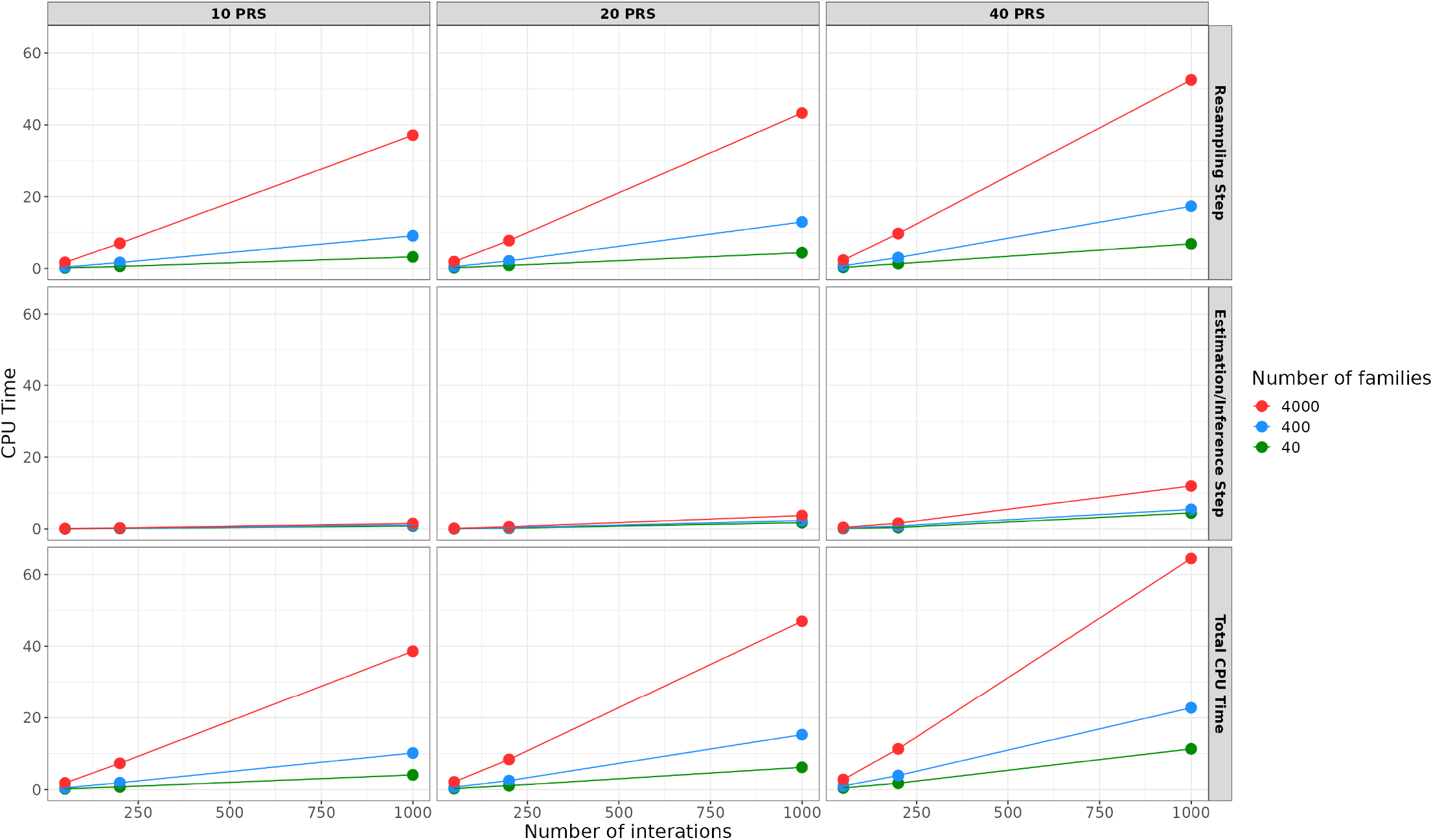
The CPU time (in seconds) of the Bootstrap algorithm over varying number of families, PRS and iterations.

## Discussion

In this study, we introduced a novel and robust Bootstrap algorithm for learning partial correlation networks from correlated data. We showed in simulated data that this algorithm effectively controls the False Positive Rate (FPR) while maintaining comparable power performance to conventional methods. Although we described the method in the context of family data, the Bootstrap algorithm can be directly applied to any correlated data setting without explicitly modeling the correlation structure, as it is required in (Ribeiro & Maria Pavan Soler, 2020).

Based on our simulation results, we recommend utilizing the Bootstrap algorithm with a minimum of 50 resampling iterations and resampling 100% of the clusters at each iteration. The algorithm should be used when there is a sufficient number of families/ clusters, such as more than 40, and large number of subjects. When sample sizes are as small as 200, resampling 100% of the families failed to control the FPR likely due to the deviation of the test statistics form a normal distribution. The Bootstrap algorithm would not be suitable, for example, in cases involving only 10 large families each with 200 relatives. Even though the total sample size is large, the limited number of clusters may not be sufficient for the Bootstrap algorithm to behave stably. Another limitation arises when the sample sizes are smaller than the number of variables, as the test statistic is no longer well-defined. When the heritability of traits is less than 25%, the Bootstrap algorithm behaves similarly to the Fisher’s test that ignores the correlation. However, in omics data analyses, most traits exhibit high heritability, typically exceeding 30%. In such cases, it would be optimal to use the Bootstrap algorithm as long as investigators have sufficient computational capacity.

To evaluate our approach, we conducted a comprehensive simulation study using genetically independent traits. It is not straightforward to extend our simulations to genetically correlated traits since the variance-covariance matrix *var*(*Y*) = (*Φ* − *I*) ⊗ *HΣ* + *I* ⊗ *Σ* is not guaranteed to be semi-positive definite when *H* is not diagonal. However, the application to the PRS in LLFS showed that our Bootstrap method works well even with some genetic correlations among traits. In fact, the heritability of PRS is very high as shown In Supplemental Table 2 and the PRS are genetically correlated since many of the outcomes shared common SNPs. In addition, we limited our simulation study to 2-generation families but it will be interesting to expand this study to multi-generation families with a variety of relatedness patterns. Finally, our simulation assumed no inbreeding and an additive genetic model, and some evaluation would be necessary to evaluate the validity of this approach to different genetic models and other types of correlated data.

The computational efficiency of our algorithm is a function of various factors, including the number of bootstrap iterations, the number of vertices, the number of families/clusters and total number of samples. The number of families impacts the resampling procedure’s runtime, while the number of nodes influences the calculation of partial correlation matrices. Furthermore, the cumulative effect of resampling and partial correlation calculations per iteration significantly contributes to the time needed for constructing Z-scores. A potential improvement to computational efficiency would involve a faster algorithm for calculating the inverse of the variance covariance matrix especially when the number of vertices are very large.

The learning strategy implemented through our proposed algorithms relies on testing multiple null hypotheses ρ_*ij*_ = 0 against the alternative hypotheses ρ_*ij*_ ≠ 0. It is important to adjust the significance levels for these tests to control the FWER. However, due to the non-independent nature of the performed tests, it is challenging to achieve precise adjustments (Ribeiro & Maria Pavan Soler, 2020). In the learning of the PRS networks, we applied the stringent Bonferroni correction to control the FWER without accounting for the effective numbers of tests, which could lead to overcorrecting as shown by (Drton & Perlman, 2007). In future work, we would like to introduce a better way to control the FWER. As an alternative to controlling the FWER, the FDR procedure by (Liu, 2013) is also a good solution that that can be integrated into our algorithm.

## Conclusion

By displaying conditional dependencies into patterns of edges in a network, GGMs offer a great statistical tool to represent intricate relationships within data in an intuitive manner and could be potentially very useful in the emerging field on multi-omics integration. However, the generation of GGMs from correlated data is a challenging task. We provided a simple method to derive a GGM from correlated data that is computationally efficient and appears to control the FPR without losing statistical power. This approach could increase the use of GGMs in observational study data that often, by design, generate correlated observations.

