## Supplemental Figure 1 for "Learning Gaussian Graphical Models from Correlated Data"

Estimates of the partial correlation at varying heritability (0-0.95) and numbers of families for $E_{i,j}$ in the triangle tail graph simulated using Bootstrap and Fisher's methods. All estimates are unbiased when the heritability is below 0.5. However, for higher heritability values, the estimates exhibit light deviations.

### Supplemental Figure 1

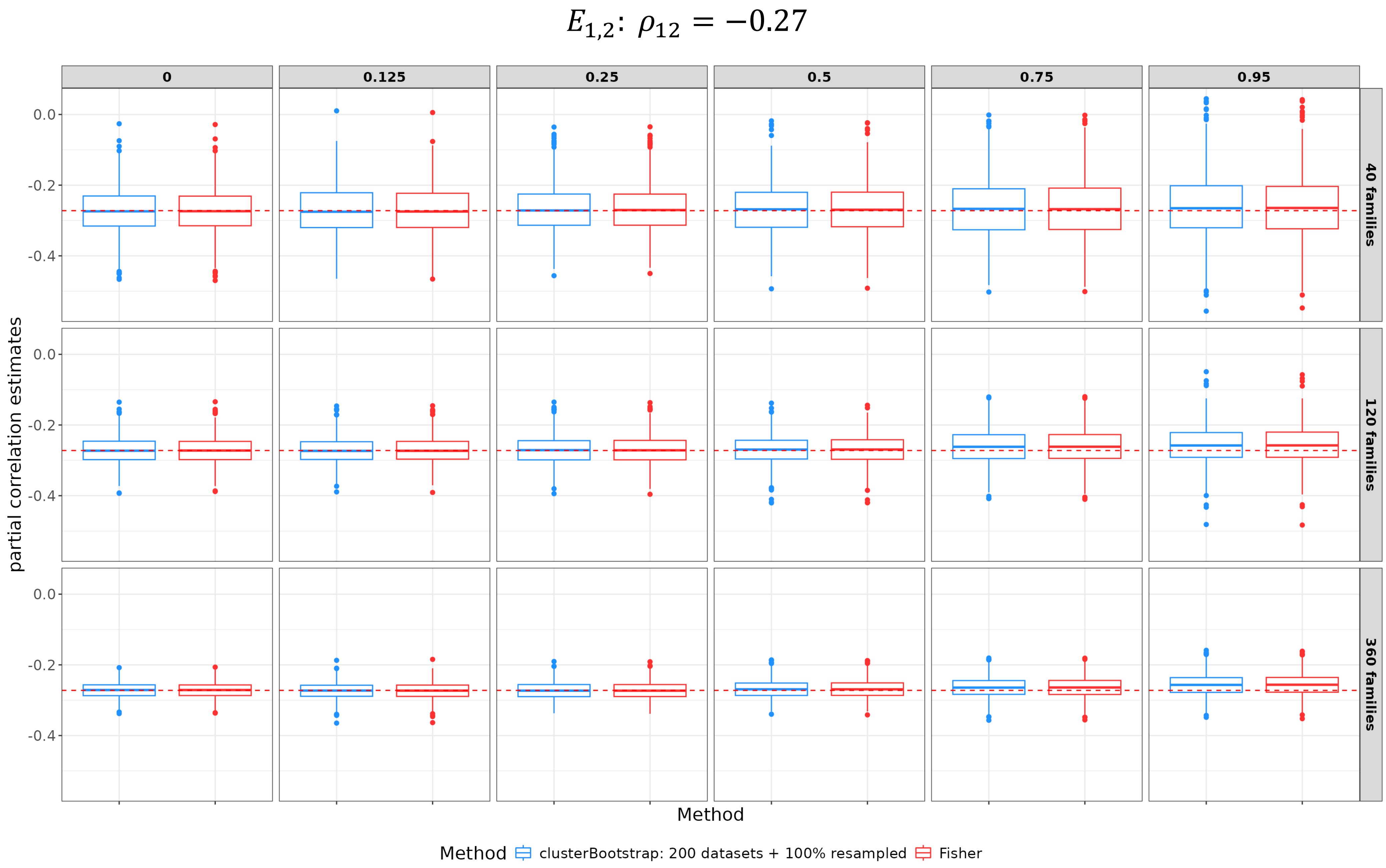

### Supplemental Figure 1

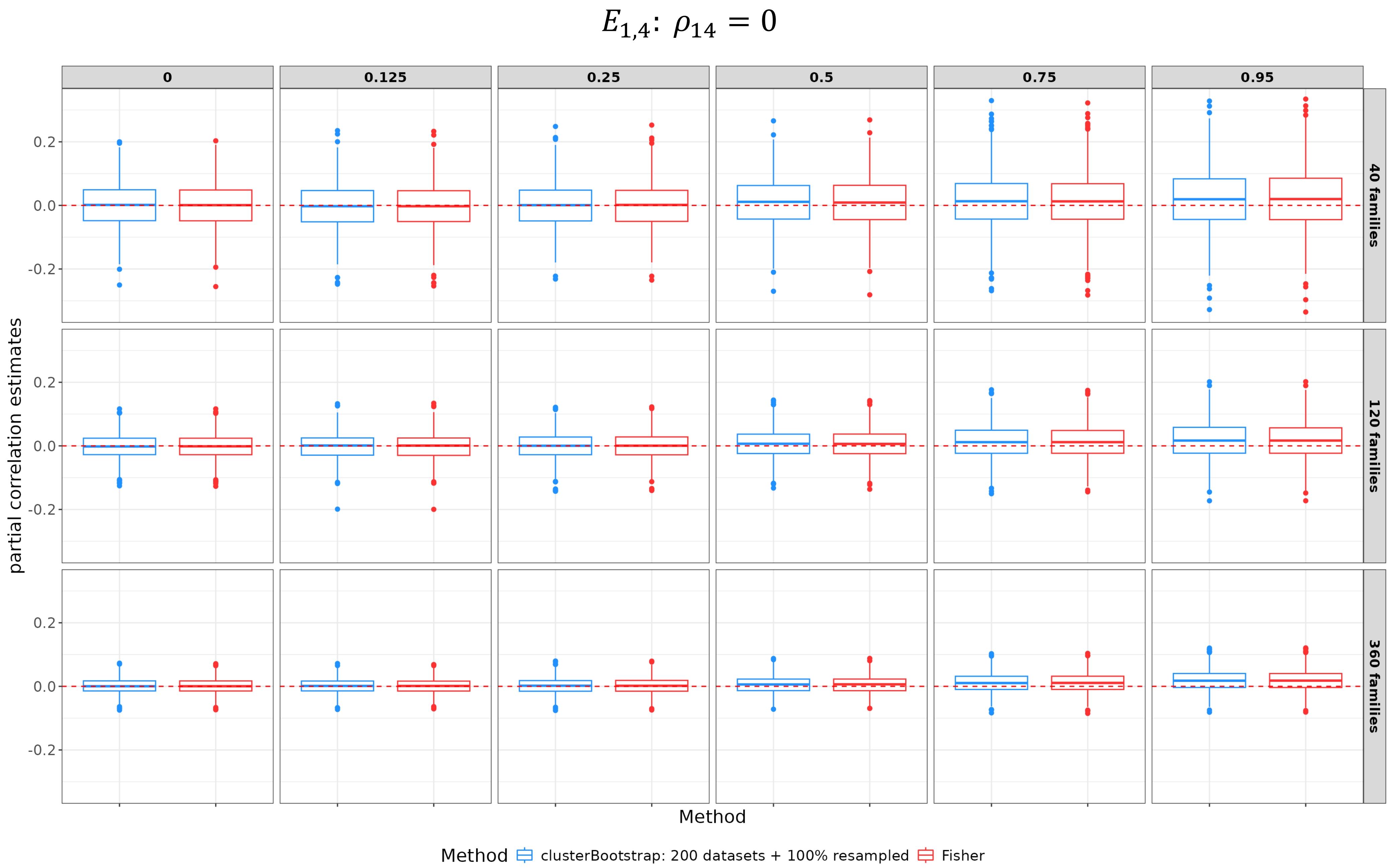

#### Supplemental Figure 1

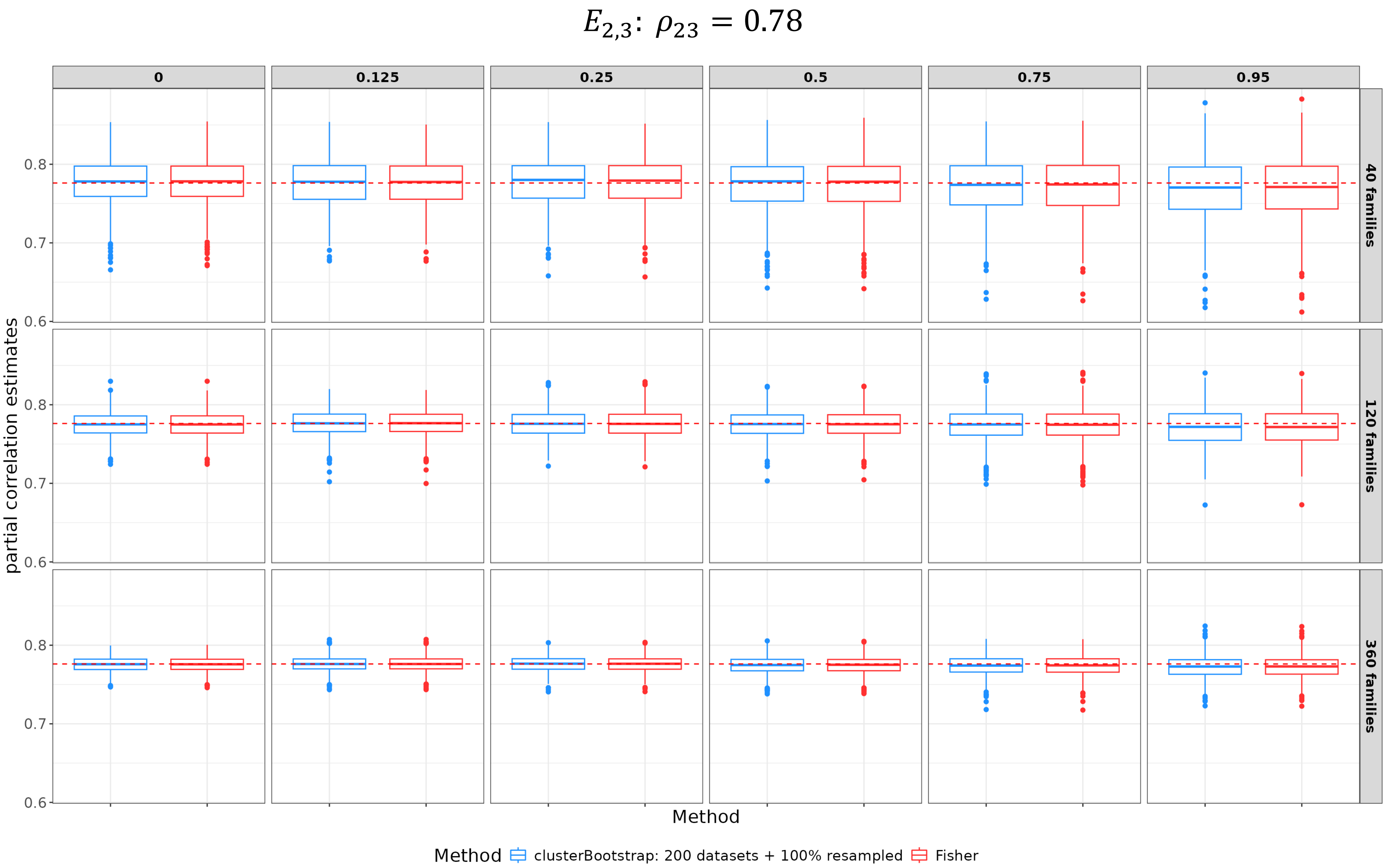

#### Supplemental Figure 1

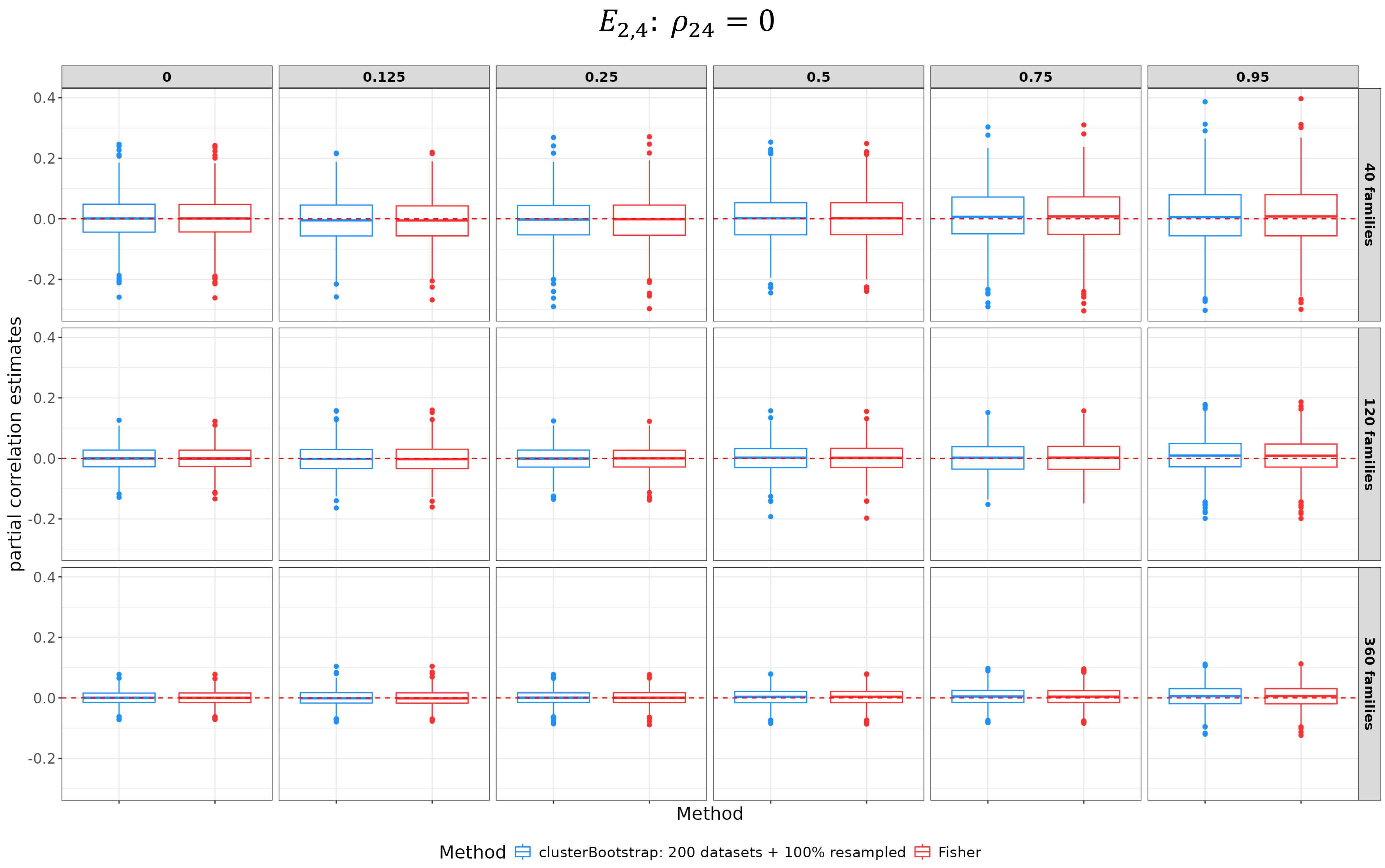

#### Supplemental Figure 1

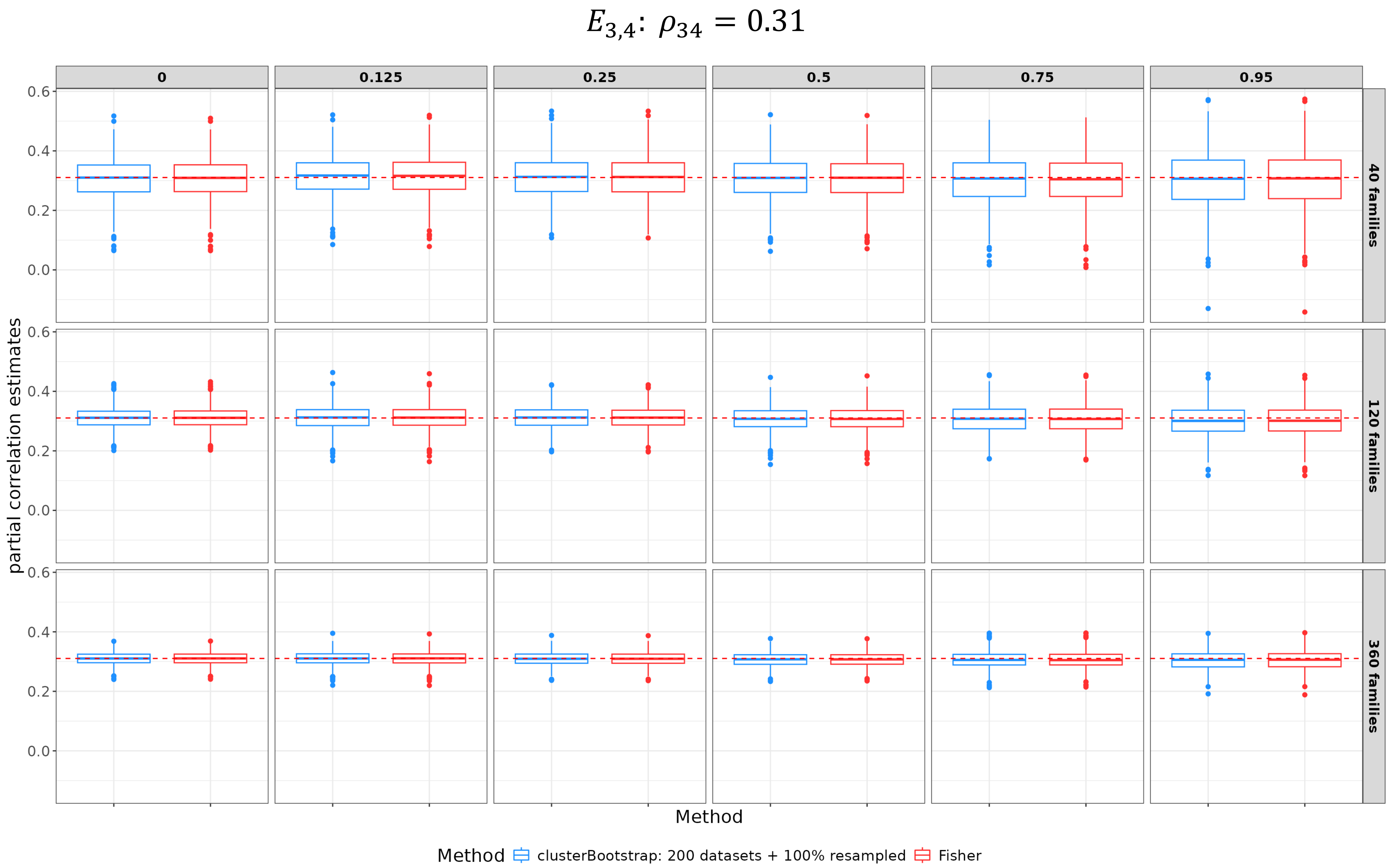
