## Supplemental Figure 2 for "Learning Gaussian Graphical Models from Correlated Data"

The histograms of 54 PRS showed that 2 PRS are not normally distributed visually.

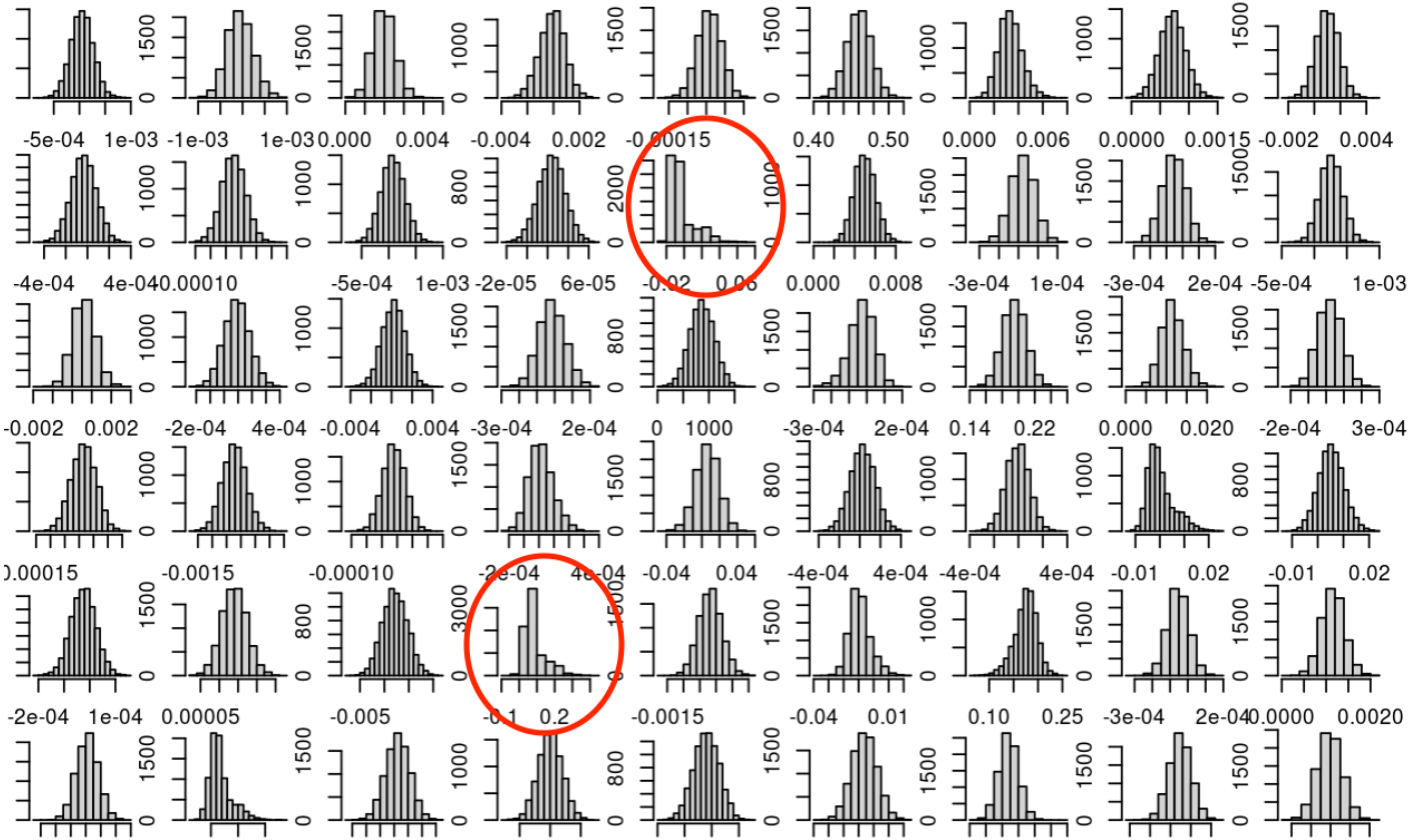
