## Supplemental Figure 3 for "Learning Gaussian Graphical Models from Correlated Data"

Scree plot of the numbers of potential outliers showed that there is a gap above 5 outliers, indicating the 5 PRS with 5 more outliers may shift away from normal distribution.

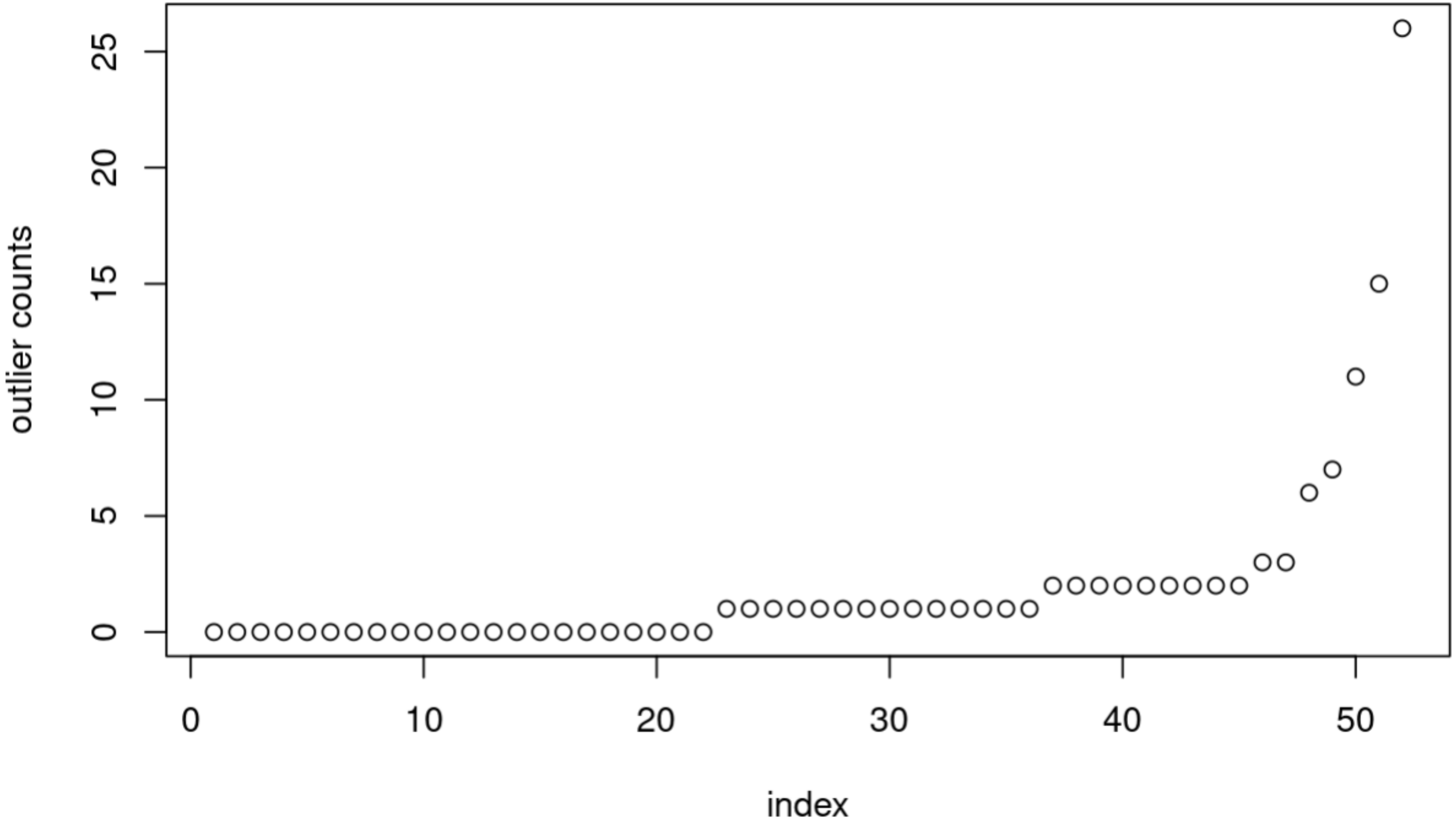
