## Supplemental Method for "Learning Gaussian Graphical Models from Correlated Data"

**Derivation of Equation 4**

Start with the generating model for a single trait $Y$:

$$Y = \mu + E + G$$

where $Y \sim N(\mu, \sigma^2)$, $E \sim N(0, \sigma_e^2)$ and $G \sim N(0, \sigma_g^2)$ and $\sigma^2 = \sigma_e^2 + \sigma_g^2$. $E$ and $G$ are assumed to be independent.

Denote the trait $Y$ as $Y_f$, $Y_m$ and $Y_o$ for father, mother, and offspring respectively. Similarly denote $E$ and $G$ as $E_f$, $E_m$ and $E_o$, and $G_f$, $G_m$ and $G_o$:

$$Y_f = \mu + E_f + G_f$$
$$Y_m = \mu + E_m + G_m$$
$$Y_o = \mu + E_o + G_o$$

By assuming $E_f$, $E_m$, and $E_o$ are independent of each other, the covariance of $Y_f$ and $Y_m$ is 0 since the parents are assumed not to be genetically correlated. The parent-offspring pair is a 1st degree relative pair and the genetic covariance between parent and offspring is: $cov(G_f, G_o) = cov(G_f, G_o) = \frac{1}{2}\sigma_g^2$.

Denote $\boldsymbol{Y} = (Y_f, Y_m, Y_o)^{\boldsymbol{T}}$, we can write the distribution of $\boldsymbol{Y}$ as:

$$\boldsymbol{Y} = \begin{bmatrix} Y_f \\ Y_m \\ Y_o \end{bmatrix} \sim MVN\left(\boldsymbol{\mu_Y}, \begin{bmatrix} \sigma^2 & 0 & \frac{1}{2}\sigma_g^2 \\ 0 & \sigma^2 & \frac{1}{2}\sigma_g^2 \\ \frac{1}{2}\sigma_g^2 & \frac{1}{2}\sigma_g^2 & \sigma^2 \end{bmatrix}\right)$$

By the theorem of conditional distribution of Multivariate Normal Distribution (MVN):

$$\begin{bmatrix} \underline{X_i} \\ \underline{X_j} \end{bmatrix} \sim MVN\left(\begin{bmatrix} \mu_i \\ \mu_j \end{bmatrix}, \begin{bmatrix} \Sigma_{ii} & \Sigma_{ij} \\ \Sigma_{ji} & \Sigma_{jj} \end{bmatrix}\right) \Rightarrow \underline{X_j} | \underline{X_i} \sim MVN\left(\mu_j + \Sigma_{ji}\Sigma_{ii}{}^{-1}\left(\underline{X_i} - \mu_i\right), \Sigma_{jj} - \Sigma_{ji}\Sigma_{ii}{}^{-1}\Sigma_{ij}\right)$$

We can derive the conditional distribution of $Y_o$ given $Y_f$ and $Y_m$ as:

$$\begin{aligned} E\left(Y_o | Y_f, Y_m\right) &= \mu + \begin{bmatrix} \frac{1}{2}\sigma_g^2 & \frac{1}{2}\sigma_g^2 \end{bmatrix} \begin{bmatrix} \sigma^2 & 0 \\ 0 & \sigma^2 \end{bmatrix}^{-1} \left(\begin{bmatrix} Y_f \\ Y_m \end{bmatrix} - \begin{bmatrix} \mu \\ \mu \end{bmatrix}\right) \\ &= \mu + \frac{1}{2}\sigma_g^2 [1 \quad 1](\sigma^2)^{-1} \begin{bmatrix} 1 & 0 \\ 0 & 1 \end{bmatrix} \begin{bmatrix} Y_f - \mu \\ Y_m - \mu \end{bmatrix} \\ &= \mu + \frac{\sigma_g^2}{2\sigma^2}\left(Y_f - \mu + Y_m - \mu\right) \\ &= \mu + \frac{\sigma_g^2}{\sigma^2}\left(\frac{Y_f + Y_m}{2} - \mu\right) \end{aligned}$$

$$\begin{aligned} var\left(Y_o | Y_f, Y_m\right) &= \sigma^2 - \begin{bmatrix} \frac{1}{2}\sigma_g^2 & \frac{1}{2}\sigma_g^2 \end{bmatrix} \begin{bmatrix} \sigma^2 & 0 \\ 0 & \sigma^2 \end{bmatrix}^{-1} \begin{bmatrix} \frac{1}{2}\sigma_g^2 \\ \frac{1}{2}\sigma_g^2 \end{bmatrix} \\ &= \sigma^2 - \frac{\sigma_g^2}{2\sigma^2}\left(\frac{1}{2}\sigma_g^2 + \frac{1}{2}\sigma_g^2\right) \\ &= \sigma^2\left(1 - \frac{1}{2}\left(\frac{\sigma_g^2}{\sigma^2}\right)^2\right) \end{aligned}$$

We thereby derived Equation 4.

**Derivation of Equation 5**

We decomposed sigma as + for simplification. Let $\underline{Y}$ be a p-dimensional Gaussian random vector $\underline{Y} = \left(Y_1, Y_2, \ldots, Y_p\right)^T$ and:

$$\underline{Y} = \boldsymbol{\mu} + \underline{E} + \underline{G}$$

where $\underline{Y} \sim N(\boldsymbol{\mu}, \Sigma)$, $\underline{E} \sim N(0, \Sigma_e)$ and $\underline{G} \sim N\left(0, \Sigma_g\right)$, and $\Sigma = \Sigma_e + \Sigma_g$.

Denote the multivariate trait $\underline{Y} = \left(Y_1, Y_2, \ldots, Y_p\right)^T$ as $\underline{Y_f} = \left(Y_{f1}, Y_{f2}, \ldots, Y_{fp}\right)^T$, $\underline{Y_m} = \left(Y_{m1}, Y_{m2}, \ldots, Y_{mp}\right)^T$ and $\underline{Y_o} = \left(Y_{o1}, Y_{o2}, \ldots, Y_{op}\right)^T$ for father, mother, and offspring respectively. Similarly denote $\underline{E}$ and $\underline{G}$ as $\underline{E_f}$, $\underline{E_m}$ and $\underline{E_o}$, and $\underline{G_f}$, $\underline{G_m}$ and $\underline{G_o}$:

$$\begin{aligned} \underline{Y_f} &= \boldsymbol{\mu} + \underline{E_f} + \underline{G_f} \\ \underline{Y_m} &= \boldsymbol{\mu} + \underline{E_m} + \underline{G_m} \\ \underline{Y_o} &= \boldsymbol{\mu} + \underline{E_o} + \underline{G_o} \end{aligned}$$

By the same reasoning as the single trait case, we evaluate the covariance of $\underline{Y_f}$ and $\underline{Y_m}$:

$$\begin{aligned} cov\left(\underline{Y_f}, \underline{Y_m}\right) &= cov\left(\underline{E_f} + \underline{G_f}, \underline{E_m} + \underline{G_m}\right) \\ &= cov\left(\underline{E_f}, \underline{E_m}\right) + cov\left(\underline{E_f}, \underline{G_m}\right) + cov\left(\underline{G_f}, \underline{E_m}\right) + cov\left(\underline{G_f}, \underline{G_m}\right) \\ &= \mathbf{0} + \mathbf{0} + \mathbf{0} + \mathbf{0} \\ &= \mathbf{0} \end{aligned}$$

Evaluate the covariances of $\underline{Y_f}$ and $\underline{Y_0}$:

$$cov\left(\underline{Y_f}, \underline{Y_o}\right) = cov\left(\underline{E_f} + \underline{G_f}, \underline{E_o} + \underline{G_o}\right)$$

$$= cov(\underline{E_f}, \underline{E_o}) + cov(\underline{E_f}, \underline{G_o}) + cov(\underline{G_f}, \underline{E_o}) + cov(\underline{G_f}, \underline{G_o})$$
$$= \mathbf{0} + \mathbf{0} + \mathbf{0} + \frac{1}{2}\Sigma_g$$
$$= \frac{1}{2}\Sigma_g$$

Denote $\boldsymbol{Y} = \left(\underline{Y_f}^T, \underline{Y_m}^T, \underline{Y_o}^T\right)^{\boldsymbol{T}}$:

$$\boldsymbol{Y} = \begin{bmatrix} Y_{f1} \\ \vdots \\ Y_{fp} \\ Y_{m1} \\ \vdots \\ Y_{mp} \\ Y_{o1} \\ \vdots \\ Y_{op} \end{bmatrix} \sim MVN(\boldsymbol{\mu_Y}, \begin{bmatrix} \Sigma & \mathbf{0} & \frac{1}{2}\Sigma_g \\ \mathbf{0} & \Sigma & \frac{1}{2}\Sigma_g \\ \frac{1}{2}\Sigma_g & \frac{1}{2}\Sigma_g & \Sigma \end{bmatrix})$$

By the theorem of conditional distribution of MVN, we can derive the conditional distribution of $\underline{Y_o}$ given their parents as:

$$E\left(\underline{Y_o}|\underline{Y_f}, \underline{Y_m}\right) = \boldsymbol{\mu} + \begin{bmatrix} \frac{1}{2}\Sigma_g & \frac{1}{2}\Sigma_g \end{bmatrix} \begin{bmatrix} \Sigma & 0 \\ 0 & \Sigma \end{bmatrix}^{-1} \left( \begin{bmatrix} \underline{Y_f} \\ \underline{Y_m} \end{bmatrix} - \begin{bmatrix} \boldsymbol{\mu} \\ \boldsymbol{\mu} \end{bmatrix} \right)$$
$$= \boldsymbol{\mu} + \frac{1}{2}\Sigma_g[I_p \quad I_p]\Sigma^{-1} I \left( \begin{bmatrix} \underline{Y_f} \\ \underline{Y_m} \end{bmatrix} - \begin{bmatrix} \boldsymbol{\mu} \\ \boldsymbol{\mu} \end{bmatrix} \right)$$
$$= \boldsymbol{\mu} + \frac{1}{2}\Sigma_g\Sigma^{-1}[I_p \quad I_p] \begin{bmatrix} \underline{Y_f} - \boldsymbol{\mu} \\ \underline{Y_m} - \boldsymbol{\mu} \end{bmatrix}$$
$$= \boldsymbol{\mu} + \Sigma_g\Sigma^{-1} \left( \frac{\underline{Y_f} + \underline{Y_m}}{2} - \boldsymbol{\mu} \right)$$

$$var\left(\underline{Y_o}|\underline{Y_f}, \underline{Y_m}\right) = \Sigma - \begin{bmatrix} \frac{1}{2}\Sigma_g & \frac{1}{2}\Sigma_g \end{bmatrix} \begin{bmatrix} \Sigma & 0 \\ 0 & \Sigma \end{bmatrix}^{-1} \begin{bmatrix} \frac{1}{2}\Sigma_g \\ \frac{1}{2}\Sigma_g \end{bmatrix}$$
$$= \Sigma - \frac{1}{2}\Sigma_g\Sigma^{-1}[I_p \quad I_p] \begin{bmatrix} \frac{1}{2}\Sigma_g \\ \frac{1}{2}\Sigma_g \end{bmatrix}$$
$$= \Sigma - \frac{1}{2}\Sigma_g\Sigma^{-1}\left(\frac{1}{2}\Sigma_g + \frac{1}{2}\Sigma_g\right)$$
$$= \Sigma - \frac{1}{2}\Sigma_g\Sigma^{-1}\Sigma_g\Sigma^{-1}\,\Sigma$$
$$= \left(I - \frac{1}{2}\left(\Sigma_g\Sigma^{-1}\right)^2\right)\Sigma$$

We thereby derived Equation 5.
