## Supplemental Table 1 for "Learning Gaussian Graphical Models from Correlated Data"

Listing of the 47 PRS and their corresponding numbers in **Figure 6.**

| Nodes | PRS |
|---|---|
| 1 | Allergic_disease |
| 2 | Alzheimers_disease |
| 3 | Amyotrophic_lateral_sclerosis |
| 4 | Ankylosing_spondylitis |
| 5 | Anxiety_tension |
| 6 | Asthma |
| 7 | Atrial_fibrillation |
| 8 | Bipolar_disorder |
| 9 | Birth_weight |
| 10 | Body_mass_index |
| 11 | Breast_cancer |
| 12 | Carpal_tunnel_syndrome |
| 13 | Chronic_kidney_disease |
| 14 | Chronotype |
| 15 | Cognitive_performance |
| 16 | Coronary_artery_disease |
| 17 | Crohns_disease |
| 18 | Depression |
| 19 | Diastolic_blood_pressure |
| 20 | Educational_attainment |
| 21 | Epilepsy |
| 22 | FEV1 |
| 23 | Glaucoma |
| 24 | Gout |

| Nodes | PRS |
|---|---|
| 25 | Heel_bone_mineral_density |
| 26 | Height |
| 27 | Inflammatory_bowel_disease |
| 28 | Insomnia_symptoms |
| 29 | Intelligence |
| 30 | Juvenile_idiopathic_arthritis |
| 31 | Life_satisfaction |
| 32 | Male_pattern_baldness |
| 33 | Narcolepsy |
| 34 | Neuroticism |
| 35 | Parental_extreme_longevity |
| 36 | Primary_biliary_cholangitis |
| 37 | Prostate_cancer |
| 38 | Sleep_duration |
| 39 | Stroke |
| 40 | Subjective_well_being |
| 41 | Systolic_blood_pressure |
| 42 | Total_body_bone_mineral_density |
| 43 | Type_1_diabetes |
| 44 | Type_2_diabetes |
| 45 | Ulcerative_colitis |
| 46 | WHRadjBMI |
| 47 | Worry |
