## Supplemental Table 2 for "Learning Gaussian Graphical Models from Correlated Data"

Top 10 heritability of PRS estimated using the linear mixed effect model

$$Y = \boldsymbol{\mu} + E + G$$

where $E \sim MVN(0, \sigma_e^2 \boldsymbol{I}_{\boldsymbol{n}\times\boldsymbol{n}})$, $G \sim MVN(0, \sigma_g^2 \boldsymbol{\Phi}_{\boldsymbol{n}\times\boldsymbol{n}})$. Heritability is estimated as

$$h^2 = \frac{\hat{\sigma}_g^2}{\hat{\sigma}_e^2 + \hat{\sigma}_g^2}$$

| PRS | Heritability | Heritability Lower CI | Heritability Upper CI |
|---|---|---|---|
| Total_body_bone_mineral_density | 1.00 | 1.00 | 1.00 |
| Insomnia_symptoms | 1.00 | 1.00 | 1.00 |
| Body_mass_index | 1.00 | 0.99 | 1.00 |
| Male_pattern_baldness | 0.99 | 0.99 | 1.00 |
| Worry | 0.95 | 0.91 | 0.99 |
| Subjective_well_being | 0.92 | 0.87 | 0.96 |
| Anxiety_tension | 0.92 | 0.88 | 0.95 |
| Stroke | 0.91 | 0.87 | 0.95 |
| Allergic_disease | 0.91 | 0.87 | 0.95 |
| FEV1 | 0.91 | 0.88 | 0.94 |
